# Task-Relevant Haptic Feedback Improves Asymptotic Performance in de novo Arm Control Acquisition

**DOI:** 10.1101/2025.04.29.651298

**Authors:** Ian S. Howard, Laura Alvarez-Hidalgo

## Abstract

The human motor system can learn to control novel effectors, but the contribution of task-relevant haptic dynamics to de novo learning remains unclear. Using a bimanual robotic interface, participants learned over two days to control the shoulder and elbow angles of a virtual arm in order to achieve accurate endpoint movements via constrained handle motions. On Day 1, one group practiced a purely kinematic mapping, whereas another group received continuous haptic feedback generated by an endpoint mass. With practice, movements shifted from sequential to more coordinated control and trajectories became straighter, with reduced directional deviation during target-directed endpoint movements, particularly in the haptic-feedback group. On Day 2, both groups learned to compensate for a velocity-dependent force field. Trajectories were initially curved but straightened with practice, and washout produced after-effects. Training in the presence of task-relevant haptic dynamics was associated with more complete error reduction during force-field exposure, while maintaining robust after-effects. Exponential modeling provided no evidence for a difference in learning rate between groups but was consistent with a lower residual (asymptotic) error in the haptic-feedback condition. These benefits therefore reflected a difference in final predictive compensation rather than in the speed of adaptation. Together, these results suggest that performance in the presence of task-relevant haptic dynamics was associated with more complete predictive compensation when adapting to novel dynamics, without evidence of faster adaptation.

## Introduction

Humans can master intricate motor skills, from rudimentary tool use to the operation of complex machinery [1–4]. Nevertheless, many motor control studies have examined relatively simple adaptations to altered task dynamics or visual feedback, often interpreted as recalibrations of existing sensorimotor mappings rather than the acquisition of fundamentally new control policies. In particular, numerous motor learning studies use veridical feedback from point-to-point movement tasks, often without kinematic transformations, and focus primarily on steady-state performance. As a result, comparatively less is known about how entirely novel movement–outcome mappings are acquired de novo, particularly when both kinematic and dynamic relationships are altered simultaneously. For example, numerous studies have investigated visuomotor adaptation [5–8], whereas dynamic learning tasks have been examined either directly [9–12] or across different unimanual contexts [13–15] and bimanual contexts [16–20].

There has also been compelling research into more complex adaptations, such as nonlinear visuomotor transformations involving movement in altered visual space [21], mirror reversal compensations [22], rotational adaptations [23], and sensorimotor transformations [24,25]. These paradigms increasingly blur the distinction between adaptation and de novo learning; however, they still typically operate within pre-existing control structures, modifying visual mappings without fundamentally restructuring the underlying motor coordination demands.

More complex remapping has been investigated using finger motions to control the joint angles of a simulated planar two-link arm, demonstrating relevance for prosthetic control [26,27]. These studies have highlighted the pivotal role of visual feedback [27] in learning to control virtual objects via finger movements [28]. Yet, in many such paradigms, visual feedback remains the dominant source of task-relevant information, and the role of physically meaningful dynamics in shaping newly acquired mappings remains comparatively underexplored.

What remains unknown is how such remapping proceeds when visual feedback is insufficient and task-relevant dynamics must be learned. Skill acquisition that requires entirely new mappings, rather than incrementally adjusting existing ones, is often referred to as de novo learning [29–31]. Such paradigms shift the focus to the acquisition of novel movement–outcome mappings that cannot be expressed as simple parametric recalibrations of previously learned control policies. Accordingly, whereas adaptation tasks may primarily involve updating parameters within an existing internal model, de novo learning requires the construction of qualitatively new sensorimotor mappings that go beyond prior control structures. Empirical evidence suggests that early phases of de novo learning are characterized by increased motor variability, reflecting exploratory behavior and uncertainty about the appropriate control policy. There is also an initial reliance on cognitive strategies, followed by the gradual emergence of more stable and automatic control policies through trial-and-error learning over practice [30,31].

The contribution of task-relevant haptic dynamics to this process remains unclear. Here, we investigate de novo learning of kinematic and dynamic relationships using a virtual two-dimensional planar arm learned either with or without task-relevant haptic feedback. The arm was implemented using a bimanual robotic manipulandum. Participants controlled the simulated arm by moving two robotic handles backward and forward through two distinct one-dimensional channels. These hand movements directly controlled the angles of the shoulder and elbow joints, thereby determining the position of the end-effector. Participants were tasked with performing center-out and out-back movements, which provided a well-structured framework to quantify changes in performance and control strategies with practice.

We examined two conditions: Training either with or without an endpoint mass attached to the virtual arm’s end-effector, allowing us to assess how exposure to different dynamic environments and the resulting haptic feedback shape the acquisition of novel kinematic and dynamic mappings. The endpoint-mass manipulation was not intended to replicate the velocity-dependent forces encountered in the curl-field phase. Instead, it provided structured, task-relevant dynamic feedback during the acquisition of the novel movement-outcome mapping. This manipulation allowed us to test whether learning a new control policy in the presence of physically meaningful dynamics influences the subsequent quality and completeness of adaptation to unpredictable forces.

In both experiments, after participants became proficient at controlling the two-dimensional arm, we evaluated their adaptation to the introduction of additional novel task dynamics, specifically a velocity-dependent curl-field, thereby enabling us to probe how prior learning history influences subsequent sensorimotor adaptation. Specifically, we asked whether prior exposure to task-relevant dynamics would influence the time course of adaptation, the final level of predictive compensation achieved, or both.

## Materials and Methods

### Participants

We recruited 16 healthy, right-handed human participants from a similar age range and educational background and randomly allocated each participant to one of two experiments. All participants provided written informed consent, and the protocol was approved by the University of Plymouth’s local ethics committee; all procedures were conducted in accordance with relevant guidelines and regulations. All participants completed the Edinburgh Handedness Inventory [32] and the Quick Physical Activity Rating (QPAR) assessment [33].

Experiment 1 (Kinematic Arm Task) comprised eight participants (4 female, 4 male; mean ± SD age 20.8 ± 1.4 years; QPAR 40.9 ± 19.4).

Experiment 2 (Endpoint Mass Task) included eight participants (3 female, 5 male; mean ± SD age 21.9 ± 3.2 years; QPAR 46.3 ± 16.3).

To avoid experimental bias, participants were kept naïve to the hypotheses of the study.

### Experimental Setup

All experiments were conducted using two identical custom-built, back-drivable vBOT planar robotic manipulanda (Fig. 1A, B), which exhibit low mass at their handles [34]. Handle position was sampled with optical encoders at 1 kHz, and planar forces were updated by torque motors at the same rate. A planar virtual reality projection system was used to overlay images of targets, the end-effector cursor, and the 2D arm configuration in the plane of the vBOTs’ movements. Participants were prevented from viewing their arms directly and were seated in front of the apparatus, holding one robot handle in each hand.

**Figure 1.**
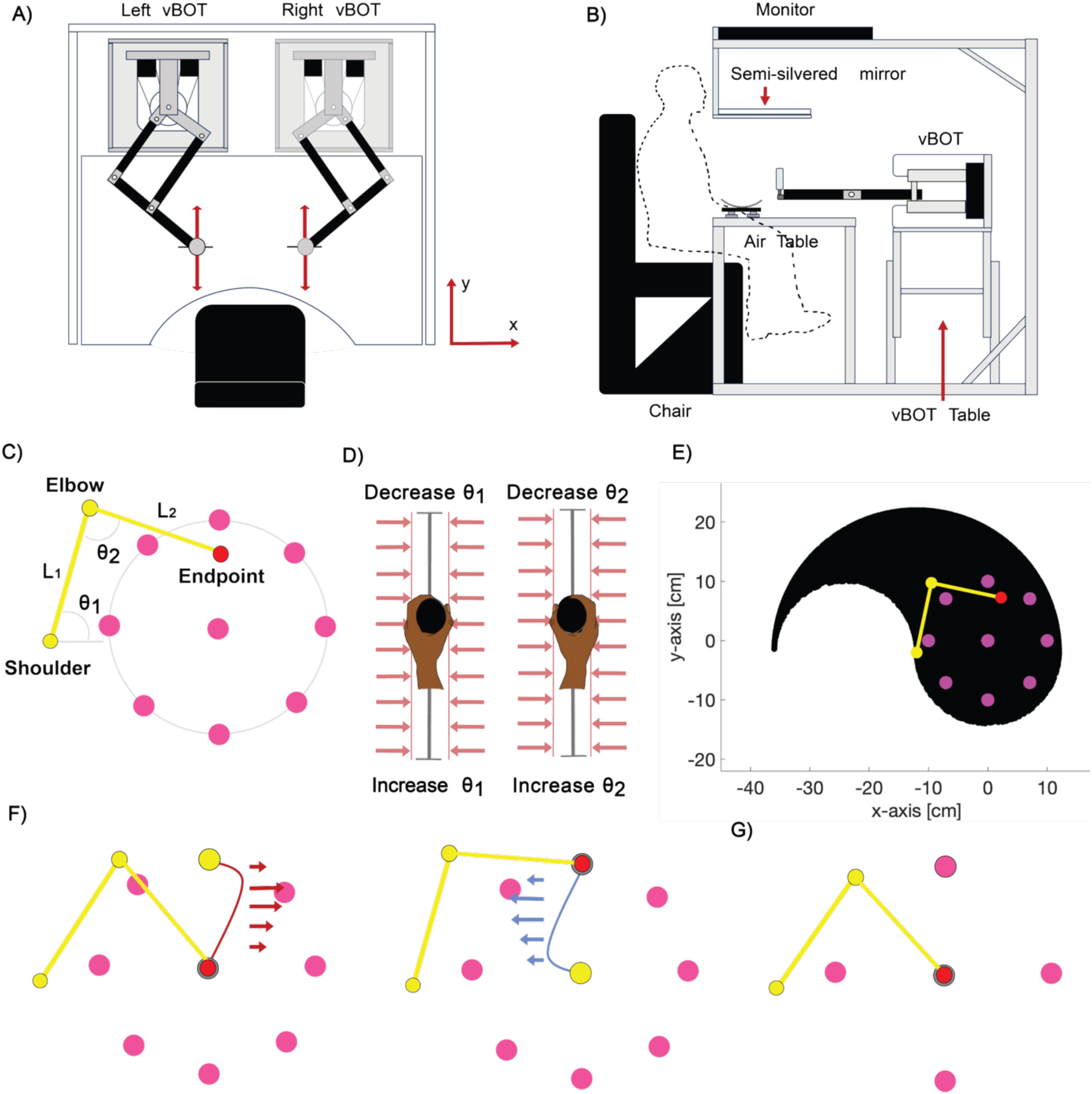
Experimental setup and task design. A: Control of a virtual two-dimensional (2D) arm using two vBOT robotic manipulanda. B: Participant operates the bimanual vBOT during the movement task while viewing the virtual display via a mirror-based 2D virtual reality system. C: Planar schematic of the simulated two-joint revolute arm. The proximal and distal links have lengths L₁ and L₂, respectively. The end-effector position is determined by joint angles θ₁ and θ₂. D: Bimanual control mapping. The left and right handles were constrained to linear channels, and their vertical positions determined the corresponding joint angles. For both joints, upward handle movement decreased the joint angle, whereas downward movement increased it. E: Reachable workspace of the virtual arm (black region). The home position and all center-out and out-back targets were located within this workspace. F: Curl-field trials. Following null-field training, participants performed outward and inward movements under velocity-dependent curl fields. Half of the participants experienced clockwise (CW) fields, and half experienced counterclockwise (CCW) fields, counterbalanced across movement directions. G: Catch trials. On these trials, movements were performed toward a single target direction selected from four possible locations (0°, 90°, 180°, or 270°).

### Two-joint Revolute Arm Simulation

We modeled a planar two-joint (shoulder–elbow) revolute arm (Fig. 1C). Full implementation details are provided in the Supplementary Information.

The workspace origin was located at (0, 0) cm. The shoulder joint was positioned at (−12, −2) cm, and both arm segments were 12 cm in length. The arm was rendered schematically, with the shoulder and elbow joints depicted as circles and the end-effector represented as a red circle at the distal endpoint. The default arm configuration was purely kinematic, with no intrinsic inertial or viscous dynamics; task-specific dynamics (e.g., endpoint mass or force-field perturbations) were introduced according to the experimental condition.

Participants controlled the arm by moving the left and right vBOT handles, which drove rotations of the shoulder and elbow joints and thereby determined the end-effector position. The sagittal (y) position of each handle was mapped monotonically to rotation of the corresponding joint (left handle for the shoulder; right handle for the elbow), thereby determining the end-effector position through forward kinematics. To spatially separate the hands, the left and right channels were offset laterally by ±10 cm, yielding central handle positions at (−10, 0) cm and (+10, 0) cm, respectively. The y-range for both hands was −15 to +15 cm.

To emphasize the novelty of the control mapping, handle motion was deliberately decoupled from intuitive endpoint motion. Specifically, outward handle movements produced inward cursor motion.

### Experimental Protocol

Participants executed center-out and out-back reaching movements with the endpoint of the two-dimensional (2D) planar arm. Each center-out movement began at a central ‘home’ position (radius: 1.0 cm; pink) and ended at one of eight peripheral targets, randomly selected without replacement (radius: 1.0 cm, pink), evenly spaced 0°, 45°, 90°, 135°, 180°, 225°, 270°, and 315° on a circle of 10 cm radius centered at (0, 0). The home start position was in the mid-sagittal plane, approximately 30 cm below eye level and 30 cm anterior to the participants’ chest. Each center-out movement was followed by an out-back movement, which involved reaching back from the previous center-out target to the central home start position.

At the beginning of each trial, the manipulanda automatically positioned the handles so the endpoint of the virtual arm was at the home location. Participants were required to keep the cursor within the central starting position and maintain cursor speed below 0.1 cm/s. After a 0.5 s delay, a target appeared at a location randomly selected without replacement, the arm became visible, and a tone indicated that the participant should move out to the target.

Each block comprised 17 trials: 8 outward (center-out) and 8 inward (out-back) movements with the simulated arm visible. In addition, one outward movement was performed with the arm invisible. The outward movements constituted a typical center-out task, enabling comparison of participants’ behaviors with prior studies. The inward (out-back) movement provided further coverage of the operating space using movements in the opposite direction. A trial terminated when the cursor was within 0.5 cm of the target and cursor speed dropped below 0.01 cm/s.

An outward catch-trial movement to a randomly selected target from one of the four cardinal directions (0°, 90°, 180°, or 270°) was also included in each block of 17 trials. In this catch trial, both the arm and the endpoint cursor were invisible, allowing assessment of feedforward control. Since catch trials are made without visual indication of cursor position, trial termination occurred when cursor speed fell below 0.01 cm/s. This criterion was used because accurate movement to a catch target was difficult and was often not achieved.

Participants were instructed to move the cursor into the target as quickly as possible and to remain stationary once the target was reached. However, unlike in many centre-out paradigms, participants were not instructed to move at a prescribed speed. This design choice was intentional, as movement duration and velocity are known to be emergent properties of motor learning, reflecting evolving control strategies rather than independent task variables. Participants were required to complete a trial movement within 10 s; otherwise, the trial was aborted. During each trial, handle positions and velocities, as well as the virtual arm’s endpoint position and velocity, were recorded.

### Experiments

We conducted two independent experiments, each comprising eight different participants, to investigate the learning and control of a virtual planar (2D) arm. Each participant completed two sessions of 2,380 trials conducted on separate days (see Fig. 2). We did not implement a familiarization period in which trial data were excluded, as we aimed to minimize the overall length of the experimental session and because initial behavior itself provides meaningful insight into participant’s movement strategies and performance.

**Figure 2.**
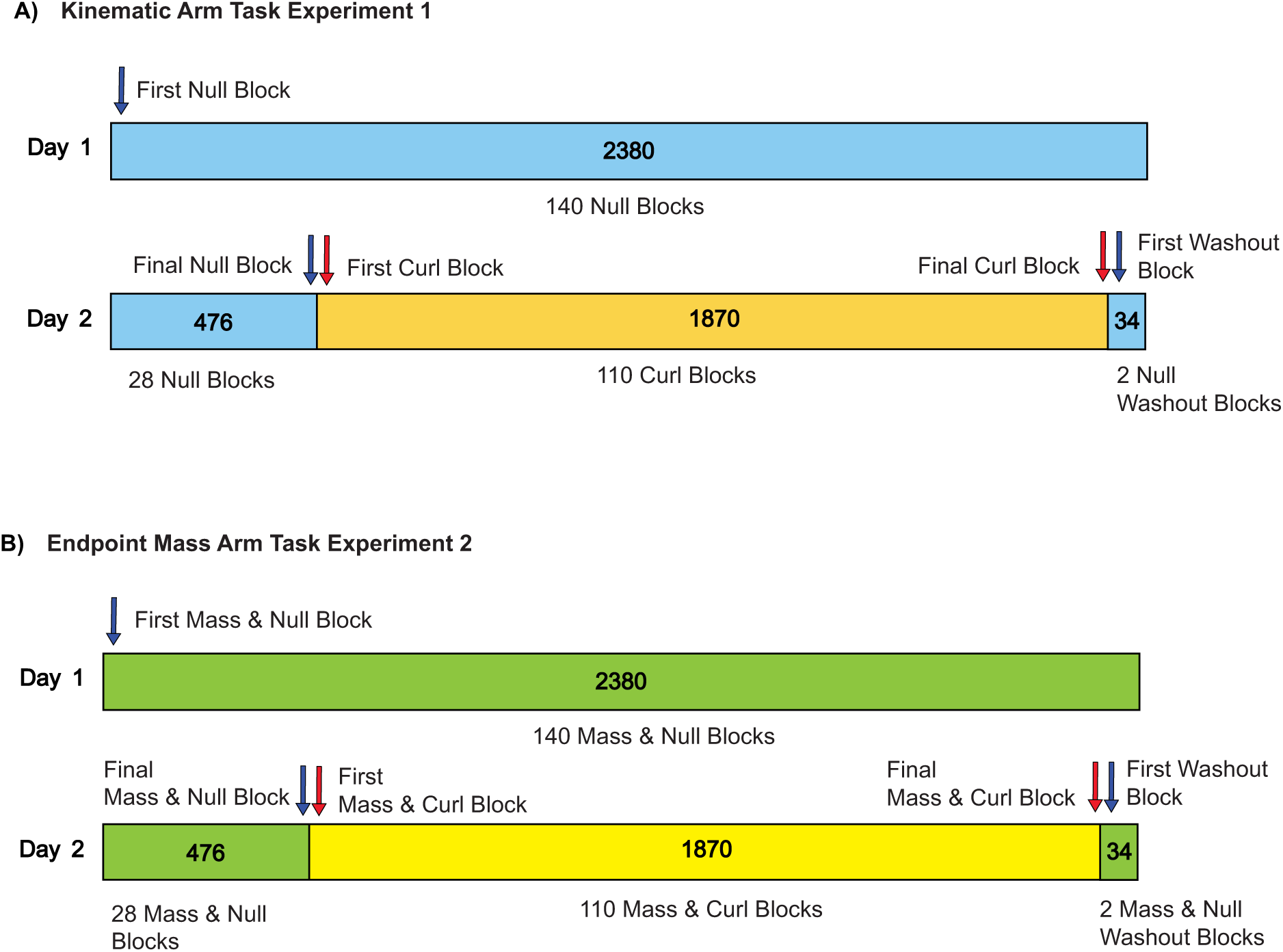
Training and testing trial schedule. A: Experiment 1. Each block consisted of 17 trials: eight cursor-visible center-out movements to a target, eight cursor-visible out-back movements to the home position, and one cursor-invisible center-out catch trial to one of four target locations. The experiment comprised null-field training, curl-field exposure, and washout phases. Trajectory analyses focused on predefined blocks: First Null, Final Null, First Curl, Final Curl, and First Washout. B: Experiment 2. Identical structure to Experiment 1, except that a simulated endpoint mass was present throughout. Analyses focused on the corresponding blocks: First Mass, Final Mass, First Mass & Curl, Final Mass & Curl, and First Mass-Only Washout.

### Kinematic Arm Task (Experiment 1)

This experiment examined de novo acquisition of kinematic control of a two-dimensional arm operating without an endpoint mass. The trial schedule is shown in Fig. 2A. On Day 1, participants executed point-to-point reaching movements in a null-field environment.

Day 2 began with a brief re-exposure to the null field, comprising the first 20% of trials (n = 476), to re-establish baseline performance. A velocity-dependent viscous curl-field was then imposed for much of the session (n = 1,870). The resulting endpoint forces were computed as joint torques via the Jacobian transpose and subsequently transformed into linear forces at the handles, enabling participants to perceive the perturbation. In the final two blocks (n = 34 trials), the curl-field was removed to assess washout after-effects following prior dynamic exposure.

### Endpoint Mass Arm Task (Experiment 2)

Experiment 2 was similar to Experiment 1, but introduced a dynamically coupled 2 kg endpoint mass, implemented via a spring–mass–damper system that generated state-dependent forces during movement. The trial schedule is shown in Fig. 2B.

In Experiment 2, the endpoint mass was present during all phases of both sessions, including null training, curl-field exposure, and washout.

The actuation forces generated by motion of the mass were first related to the virtual arm joint torques and subsequently mapped to corresponding forces at the control handles. Participants were not informed about the nature of the endpoint dynamics, and the endpoint-mass condition was designed to provide state-dependent forces related primarily to movement acceleration, rather than only velocity-dependent perturbations.

## Data Analysis

The experimental data were processed offline using MATLAB R2024a.

### Hand Trajectory Plots

Hand trajectory plots provide an intuitive visualization of participants’ movement behavior throughout the experiment. They illustrate naïve control early in training, the progressive refinement that accompanies exposure to external dynamics (e.g., viscous curl-fields), and the re-emergence of baseline patterns once those forces are removed, thereby revealing learning, adaptation, and de-adaptation within a single visualization.

In both experiments, several key phases are critical to understanding participants’ motor performance and adaptation. These include the first and final blocks of null-field trials (baseline establishment and improvement), the first and final blocks of curl-field exposure (early and late adaptation), and the first washout block (de-adaptation).

Hand trajectory plots allow direct comparison of movements during outward and inward paths and provide insight into both extrinsic trajectories (endpoint-space paths) and intrinsic trajectories (handle y-positions, which determine the corresponding joint angles). Additionally, catch trials, in which the target is shown but no visual feedback of the arm is provided, are included to assess the development of feedforward predictive control used to manipulate the novel structure of the 2D arm.

Trajectory quantification: Several measures were calculated to quantify the movement trajectory of each individual trial. First, the movement start and end points were determined in both time and space. A movement was considered to have started when the center of the extrinsic cursor left the center of the starting target circle (radius: 0.5 cm) and was traveling at a speed greater than 0.01 cm/s. A movement was considered to have stopped when the center of the extrinsic cursor reached the extrinsic target circle (radius: 0.5 cm). For catch trials, in which the cursor was invisible, movement termination was instead defined by cursor speed falling below the threshold.

### Movement Speed and Duration

Movement duration and peak speed were analyzed as dependent variables rather than controlled parameters. This approach is consistent with prior work on de novo learning, where changes in speed reflect the stabilization of control policies. To ensure that group differences in adaptation were not trivially driven by speed, we also performed analyses that normalized deviation metrics by peak movement speed (see Results).

### Performance Metrics for Visible Movements

We quantified the kinematic accuracy of visible movements with four complementary deviation metrics.

#### Extrinsic Signed Maximum Perpendicular Error (SMPE)

We calculated the signed maximum deviation of the 2D arm endpoint trajectory from a straight-line path connecting the starting cursor location and the target cursor location in extrinsic space. This measure preserves the direction of the maximum deviation from a straight-line trajectory. Because deviations retain their sign, averaging SMPE across trials yields a measure of systematic directional bias, as positive and negative deviations offset one another. In contrast, purely unsigned measures do not distinguish between bias and variability. As a result, the group-mean SMPE is a clearer indicator of systematic bias in the trajectories. Accordingly, since its introduction, this has been the preferred metric used to assess deviation from a straight line in curl-field adaptation experiments [35].

Adaptation to the curl-field was primarily assessed using SMPE, which captures systematic directional bias and indexes predictive compensation.

#### Speed-normalized curl-field analysis (SMPEN)

To ensure that differences in experienced curl-field force magnitude did not trivially account for group differences, signed mean path error (SMPE) was normalized by peak movement speed.

#### Extrinsic Absolute Maximum Perpendicular Error (AMPE)

For each trial, we took the absolute value of the maximum perpendicular deviation of the 2D arm endpoint trajectory from the ideal straight-line path joining the start cursor location and the target cursor location in extrinsic space. This measure ensures that positive and negative deviations over trials do not cancel out and gives an overall indication of how well participants were able to move along a straight trajectory from the starting point to the target, making it a suitable metric to quantify overall control of the virtual arm.

#### Movement Duration

The difference between movement onset and termination times.

#### Movement Length

To estimate the extrinsic movement path length, we computed the sum of the distances between adjacent (x, y) cursor trajectory positions, calculated using the Pythagorean theorem, from the start to the end of the movement trajectory.

### Mean ± SE over Trials and Participants

The average values of each trajectory metric were first computed across all trials within a given set of blocks. The mean ± standard error (SE) of the resulting average values was then calculated across all participants.

### Performance Metrics for Catch Movements

Catch trials were used to assess predictive control of the learned mapping in the absence of visual feedback. Because no online visual correction was possible and trials terminated based on movement cessation rather than target acquisition, catch-trial performance reflects feedforward movement planning under uncertainty about the endpoint location. Catch-trial performance was quantified using two complementary termination-based metrics.

#### Trial-end distance from start

The Euclidean distance between the start location and the endpoint at movement termination, defined by cursor speed falling below the threshold, was calculated.

#### Trial-end distance from target

The Euclidean distance between the target location and the endpoint at movement termination was calculated.

### Statistics

Statistical analyses were conducted in JASP [36]. A Mauchly’s test of sphericity was applied to check for sphericity wherever applicable. In cases where the Mauchly’s test was significant, the degrees of freedom were adjusted using a Greenhouse-Geisser correction. Parameter α was set to 0.05 in all tests. We report omega squared as the effect size (ω²).

Each of the two experiments was conducted with different participants, resulting in a total of two sets of SMPE, AMPE, path length, and movement duration data. For each of these two datasets in isolation, we first performed repeated measures ANOVAs to examine differences in SMPE, AMPE, path length, and duration data at various critical phases within each experimental condition. We also similarly analyzed SMPEN, the speed-normalized measure.

To evaluate these metrics within each experimental condition, we compared the mean value of each metric over two blocks for the initial null phase, the final null phase, the initial curl-field exposure phase, the final curl-field exposure phase, and the washout phase. This comparison was conducted using a repeated-measures ANOVA, with phase as a repeated-measures factor (five levels). When a significant main effect was observed, we proceeded with post-hoc comparisons using the Holm-Bonferroni correction.

To compare results across experiments for the metrics, we analyzed the mean of each metric over ten blocks using an ANOVA across the final null phase, the initial curl-field exposure phase, and the final curl-field exposure phase. In this analysis, experimental condition (two levels) was considered the factor. Post-hoc comparisons were conducted with the Holm-Bonferroni correction if a significant main effect was detected.

### Learning-curve analysis

To assess differences in SMPE learning curves during curl-field exposure between conditions, we fitted an exponential function to block-pair averaged SMPE data. Group-level parameter differences were evaluated using a participant-level permutation test. Participants from both conditions were pooled and repeatedly reassigned without replacement to two pseudo-groups matching the original group sizes. For each permutation, the exponential model was refitted to block-pair data within each pseudo-group and parameter differences were computed. Two-sided p-values were calculated as the proportion of permuted parameter differences whose absolute magnitude exceeded that of the observed difference. Full details are provided in the Supplementary Material.

## Results

In Experiment 1, participants first learned an arbitrary kinematic mapping between hand movements and joint angles before exposure to an endpoint curl field (Fig. 2A). Experiment 2 followed the same structure but included endpoint mass throughout training (Fig. 2B).

### Trajectory Plots Kinematic Arm Task Experiment 1

Representative trajectories illustrate the progression of de novo learning (Fig. 3A–B). Early in the null phase, extrinsic paths were erratic and intrinsic motions segmented, reflecting independent hand control. By the final null block, trajectories became straighter and intrinsic motions more continuous. Across participants (Fig. 3C–D), null-phase learning reduced curvature, curl-field introduction induced systematic deviations, and continued exposure restored straighter trajectories. Washout produced clear after-effects in the opposite direction of the imposed perturbation.

**Figure 3.**
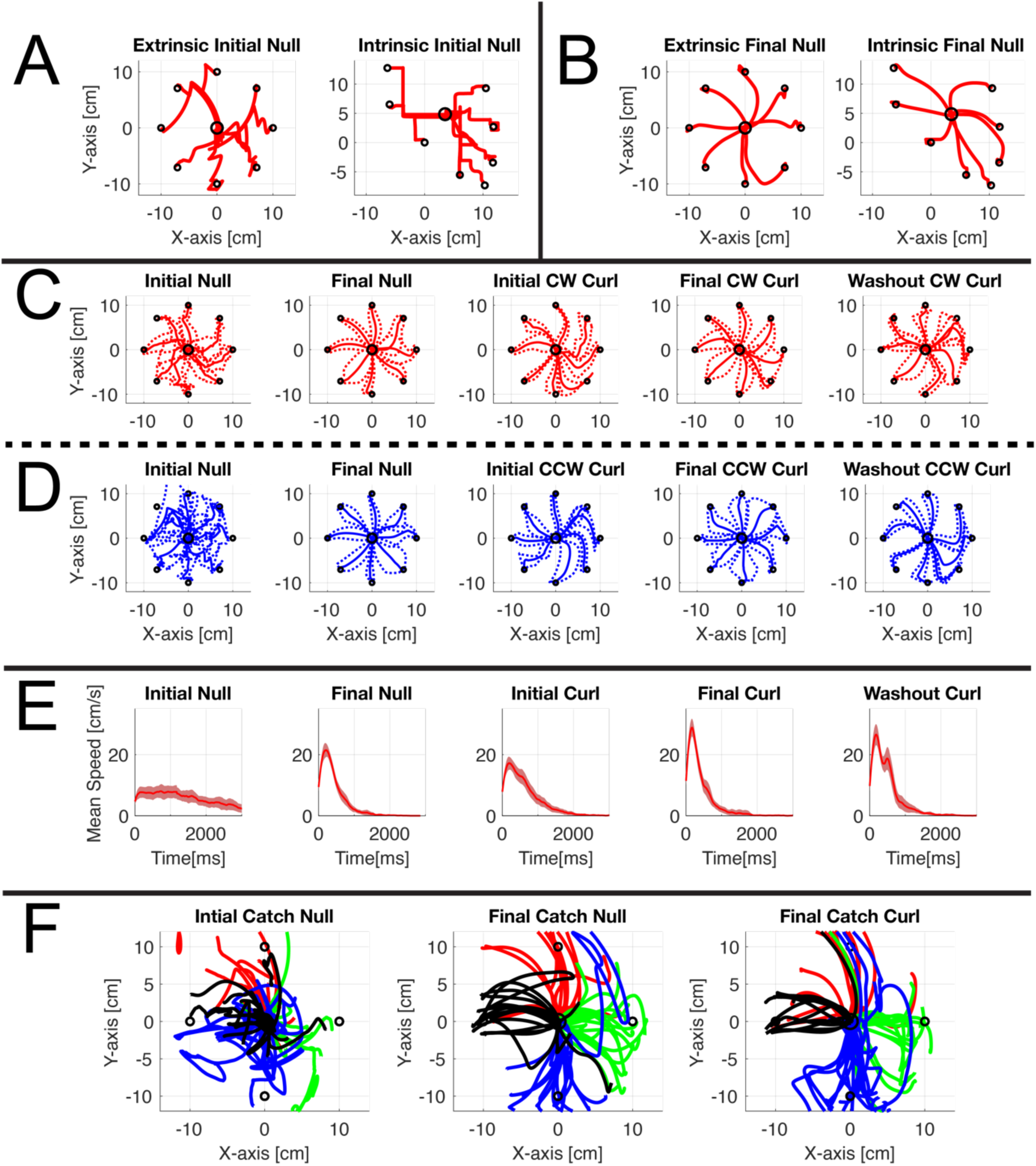
Trajectory plots for Experiment 1 – Kinematic Arm Task. **A:** Representative extrinsic (arm endpoint) and intrinsic (handle y-positions) trajectories from the first null block of an individual participant. **B:** Representative extrinsic (arm endpoint) and intrinsic trajectories (handle y-positions) from the final null block of the same participant. **C:** Mean ± SE extrinsic outward trajectory paths for the clockwise (CW) curl-field condition during the initial null block, final null block, initial curl block, final curl block, and washout blocks. **D:** Corresponding mean ± SE extrinsic outward trajectory paths for the counterclockwise (CCW) curl-field condition. **E:** Corresponding mean ± SE extrinsic speed profiles collapsed across CW and CCW trials. **F:** Extrinsic catch-trial trajectories from the first eight null blocks, the final eight null blocks, and the final eight curl blocks. Trajectories to the 0°, 90°, 180°, and 270° targets are plotted in red, green, blue, and black, respectively.

Mean speed profiles (Fig. 3E) evolved from slow, flattened shapes during early learning toward more peaked, approximately minimum-jerk-like profiles by the end of the null phase. Curl-field introduction reduced peak speed and increased asymmetry, followed by partial recovery during adaptation. Washout increased variability consistent with after-effects.

Catch trials without visual feedback (Fig. 3F) revealed progressive improvements in feedforward control. Early movements were frequently misdirected and mis-scaled. By the end of null training and curl-field exposure, directional accuracy and movement extent improved, although variability remained across participants.

### Trajectory Plots Endpoint Mass Task Experiment 2

Trajectory patterns in Experiment 2 paralleled those in Experiment 1 (Fig. 4). Early movements were slower, consistent with a stabilization strategy during acquisition of a novel control policy. Curl-field exposure again induced systematic curvature followed by recovery during adaptation. Final SMPE values approached baseline more closely than in Experiment 1.

**Figure 4.**
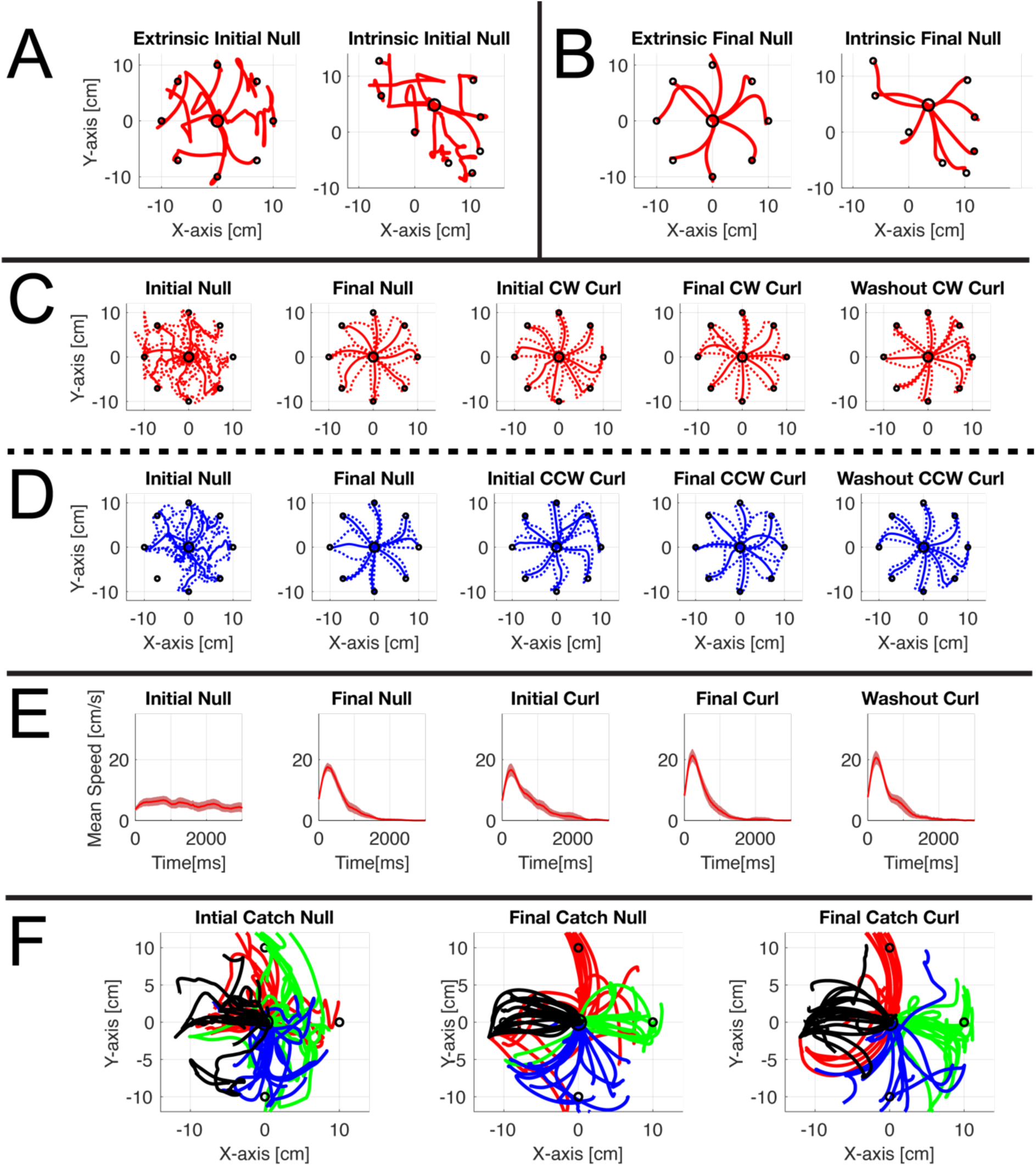
Trajectory plots for Experiment 2 – Endpoint Mass Task. **A:** Representative extrinsic (arm endpoint) and intrinsic (handle y-positions) trajectories from the first null block of an individual participant. **B:** Representative extrinsic (arm endpoint) and intrinsic (handle y-positions) trajectories from the final null block of the same participant. **C:** Mean ± SE extrinsic outward path trajectories for the CW curl-field condition. **D:** Mean ± SE extrinsic outward path trajectories for the CCW curl-field condition. **E:** Mean ± SE extrinsic speed profiles across CW and CCW trials. **F:** Extrinsic catch-trial trajectories for the first eight null blocks, the final eight null blocks, and the final eight curl blocks. Trajectories to the 0°, 90°, 180°, and 270° targets are plotted in red, green, blue, and black, respectively.

### Examining Effects Within Experiment 1

Movement trajectory quality was quantified using SMPE, AMPE, movement duration, and extrinsic path length (Fig. 5). Metrics were averaged across paired blocks (32 trials per pair). A small discontinuity is visible at the start of Session 2 due to the separation across days.

**Figure 5.**
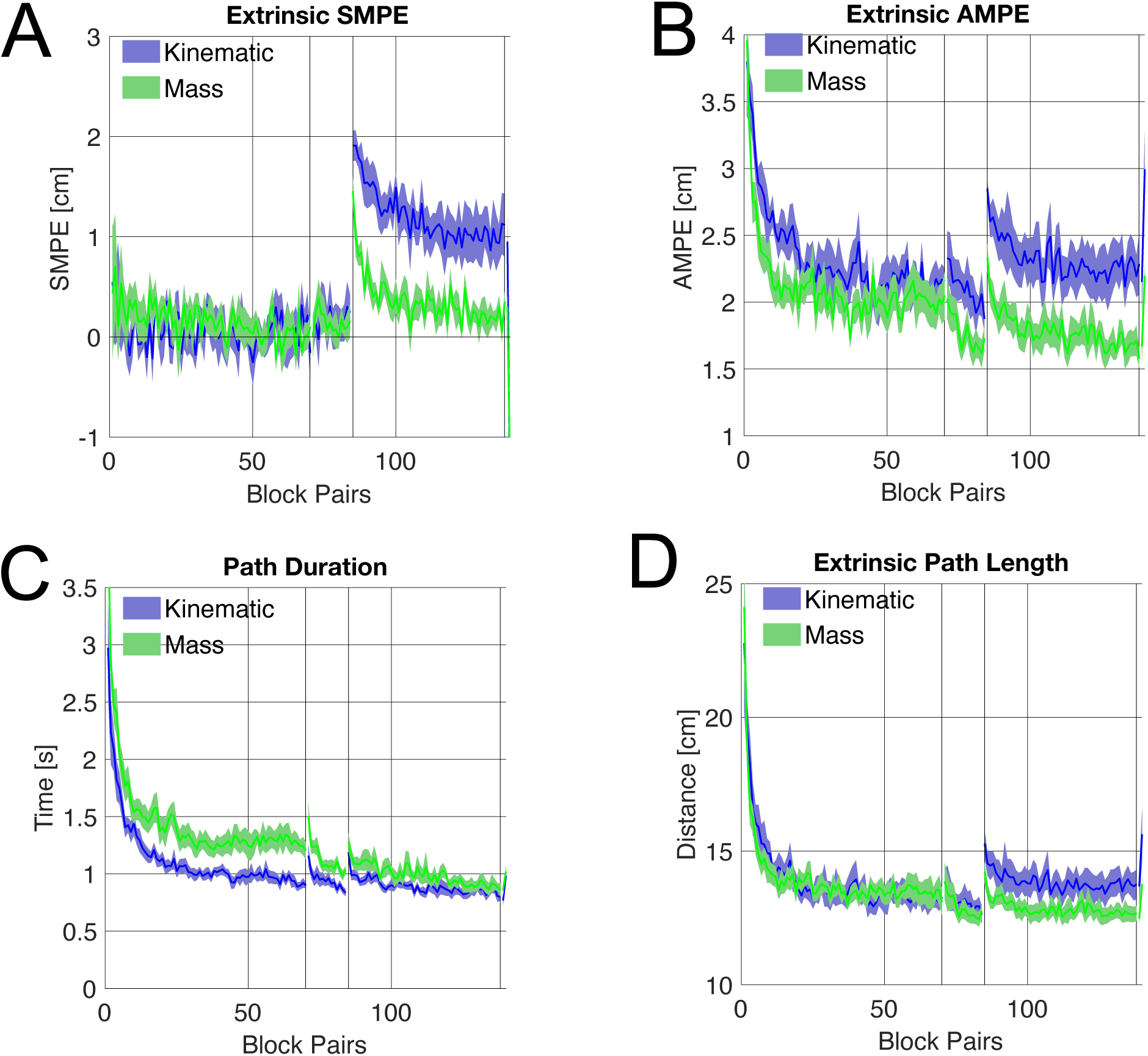
Comparison of movement metrics averaged over pairs of trial blocks (32 trials per point) between Experiment 1 – Kinematic Arm Task (blue) and Experiment 2 – Endpoint Mass Task (green). Statistical comparisons across experiments were performed at the end of null-field learning, the initial curl-field exposure phase, and the final curl-field exposure phase. **A:** Mean ± SE extrinsic signed maximum perpendicular error (SMPE) over the experiments. **B:** Mean ± SE extrinsic absolute maximum perpendicular error (AMPE) shown over the experiments. **C:** Mean ± SE trial duration over the experiments. **D:** Mean ± SE extrinsic path length over the experiments.

Within each experiment, performance metrics were averaged over 2-block pairs (32 trials) so that trial-to-trial variability was reduced without obscuring relatively rapid changes across learning phases.

To assess within-experiment changes, we compared performance across five critical stages: initial null, final null, initial curl, final curl, and washout. Full statistical results are provided in Supplementary Table S1.

### Learning in the Null Phase

During de novo arm control training (initial to final null), participants reduced unsigned error (AMPE) and movement duration, whereas signed error (SMPE) and path length showed a slight change (Table S1). This indicates improved trajectory efficiency without systematic directional bias correction prior to perturbation exposure.

### Curl-Field Introduction and Adaptation

Introduction of the curl field produced a marked increase in error and movement duration across metrics. With continued exposure, all trajectory measures showed significant recovery toward baseline levels, consistent with predictive adaptation (Fig. 5; Table S1).

### Washout and After-Effects

Following removal of the curl field, participants exhibited systematic after-effects, characterized by signed deviations opposite to the imposed perturbation. SMPE during early washout was consistent with predictive compensation rather than passive attenuation from co-contraction. Error and path metrics differed significantly from the final curl phase, whereas movement duration did not fully rebound (Table S1).

### Movement Metrics Plots Experiment 2

Experiment 2 followed the same structure as Experiment 1 but with mass present throughout training. However, asymptotic error during curl-field exposure was lower in the endpoint-mass condition. The overall pattern of null learning (Fig. 6), curl-field disruption, adaptation, and washout mirrored that of Experiment 1 (Table S2). As in Experiment 1, introduction of the curl field increased trajectory error, followed by progressive recovery during exposure. After-effects during washout again confirmed predictive compensation.

**Figure 6.**
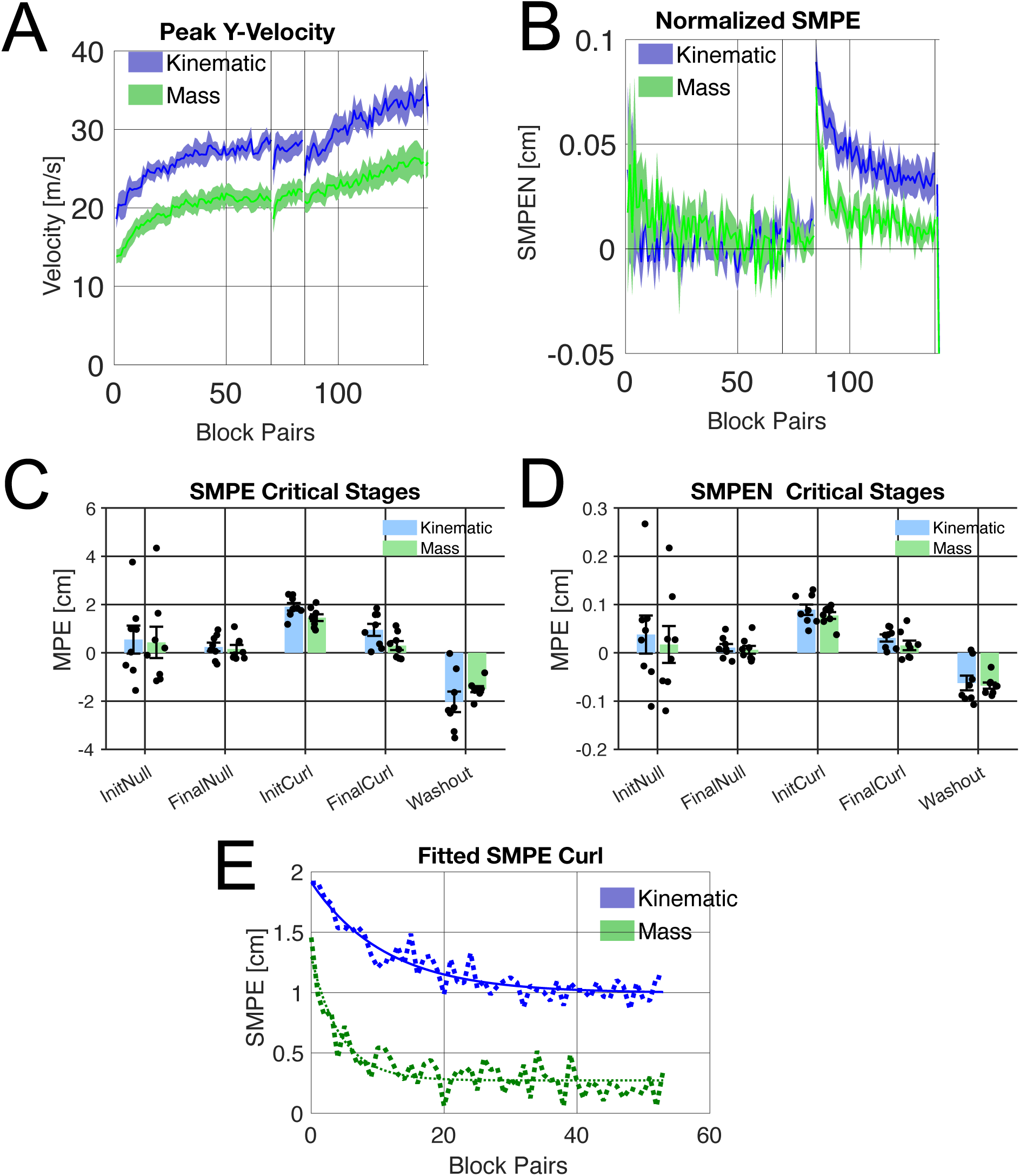
Examination of the effect of peak velocity. **A**: Peak velocity toward the target (mean ± SE). **B**: Signed maximum perpendicular error (SMPE) normalized by peak velocity. **C**: SMPE at critical stages of the experiment. **D**: Normalized SMPE (SMPEN) at critical stages. **E**: Exponential fits (solid lines) applied to pooled block-pair averaged SMPE learning curves during curl-field exposure (each data point representing the mean of 32 curl-field trials, excluding catch trials). Permutation testing provided insufficient evidence for a between-condition difference in learning rate (τ), whereas the endpoint-mass condition exhibited a significantly lower asymptotic error (C), indicating more complete predictive compensation. Panels C and D summarize values at the critical experimental stages. Panel E shows exponential fits applied over the curl-field exposure period.

### Comparison of Metrics Across Experiments

To compare the two experimental conditions, we examined SMPE, AMPE, movement duration, and path length across block pairs (Fig. 5). In addition, ANOVAs were performed on metrics averaged over 10 blocks (160 trials) at three critical stages: end of null-field learning, initial curl-field exposure, and final curl-field exposure.

Between experiments, performance metrics were averaged over 10 blocks (160 trials) to obtain more reliable estimates of performance at key stages of the experiment.

Quantitative comparisons of washout between conditions were not performed due to the limited number of washout blocks, which were considered insufficient for reliable statistical inference.

### Signed Maximum Perpendicular Error (SMPE)

During the final null phase, there was no evidence for a difference between experiments (F(1,14) = 0.176, p = 0.681, ω² < 0.001).

At curl-field onset, however, a significant between-condition difference emerged (F(1,14) = 32.405, p < 0.001, ω² = 0.662), with the endpoint-mass condition showing reduced directional bias.

This difference persisted at the end of curl-field exposure (F(1,14) = 9.015, p = 0.010, ω² = 0.334). The endpoint-mass condition exhibited lower asymptotic signed error relative to the kinematic condition.

In both conditions, early washout exhibited sign-reversed after-effects consistent with predictive compensation; between-group comparisons were not performed due to the limited number of washout blocks.

### Absolute Maximum Perpendicular Error (AMPE)

AMPE was modestly lower in the endpoint-mass condition at the end of null-field learning (F(1,14) = 4.778, p = 0.046, ω² = 0.191).

At curl-field onset (F(1,14) = 15.405, p = 0.002, ω² = 0.474) and at the end of curl-field exposure (F(1,14) = 8.492, p = 0.011, ω² = 0.319), the mass condition continued to exhibit significantly lower unsigned trajectory error.

Thus, the endpoint-mass condition exhibited reduced unsigned trajectory deviation both before and during exposure to novel dynamics.

### Movement Duration and Path Length

Movement duration was longer in the endpoint-mass condition during the null phase (F(1,14) = 18.603, p < 0.001, ω² = 0.524) and at curl onset (F(1,14) = 6.952, p = 0.020, ω² = 0.271), but not at the end of curl exposure (F(1,14) = 2.218, p = 0.159, ω² = 0.071). This difference in duration is consistent with the higher inertial load in the endpoint-mass condition.

In contrast, extrinsic path length did not provide statistically significant evidence for differences between conditions at any stage: End of null-field learning (F(1,14) = 0.088, p = 0.770, ω² = 0.000), curl onset (F(1,14) = 2.939, p = 0.109, ω² = 0.108), end of curl exposure (F(1,14) = 2.893, p = 0.111, ω² = 0.106).

Together, these results indicate that the endpoint-mass condition was associated with consistently lower signed (directional) error as indexed by SMPE, as well as lower unsigned trajectory error (AMPE), without altering overall path geometry.

### Effect of Movement Velocity

Because curl-field forces scale linearly with movement velocity, differences in trajectory error could reflect differences in movement speed. Peak velocity differed between conditions (Fig. 6A). However, after normalizing SMPE by peak velocity, the endpoint-mass condition continued to exhibit lower error during curl-field exposure (Fig. 6B).

Washout after-effects were also preserved after speed normalization, indicating predictive compensation. The between-condition difference persisted after speed normalization.

### Learning Time Constant Analysis

To determine whether between-condition differences reflected adaptation rate or asymptotic performance, we fitted an exponential model to curl-field SMPE learning curves computed over block pairs (each data point representing the mean of 32 curl-field trials, excluding catch trials; see Fig. 6E and Supplementary Information for methodological details). The time constant τ therefore reflects adaptation across block pairs rather than individual trials.

Participant-label permutation testing provided insufficient evidence for between-condition differences in learning rate (τ; p = 0.242) or initial amplitude (A; p = 0.584). However, the asymptotic parameter (C) differed significantly between conditions (p = 0.012), with the endpoint-mass condition achieving lower residual error. Parameter estimates are provided in the Supplementary Material.

Applying the same analysis to velocity-normalized SMPE (SMPEN) yielded a qualitatively similar pattern: no evidence for a between-condition difference in learning rate (τ), but a significant difference in asymptotic error (p = 0.043).

These modeling results are consistent with the late-phase SMPE comparisons, indicating that the difference between-conditions was expressed primarily in final performance rather than in adaptation rate.

### Feedforward Control Evidence from Catch Trials

Catch trials, in which visual feedback of both cursor and arm was removed, were used to assess feedforward control of the learned movement–outcome mapping. Because no online visual correction was possible and trial termination was based solely on movement cessation, catch-trial performance reflects predictive specification of movement extent rather than visually guided endpoint correction.

Initial catch-trial movements exhibited highly variable and inconsistent movement directions, particularly early in learning. Directional error measures showed high variability and were therefore not analyzed quantitatively. Accordingly, catch-trial analysis focused on movement extent at trial termination, which provides a robust indicator of feedforward control under these conditions.

Deviation from the start point at movement termination increased systematically across training (Fig. 7A), indicating that participants progressively produced movements of greater extent rather than prematurely terminating movements. Conversely, deviation from the target point at termination decreased across training (Fig. 7B). Given their low frequency (1 per 17 trials), catch trials were treated as qualitative indicators of feedforward control rather than primary inferential outcomes.

**Figure 7.**
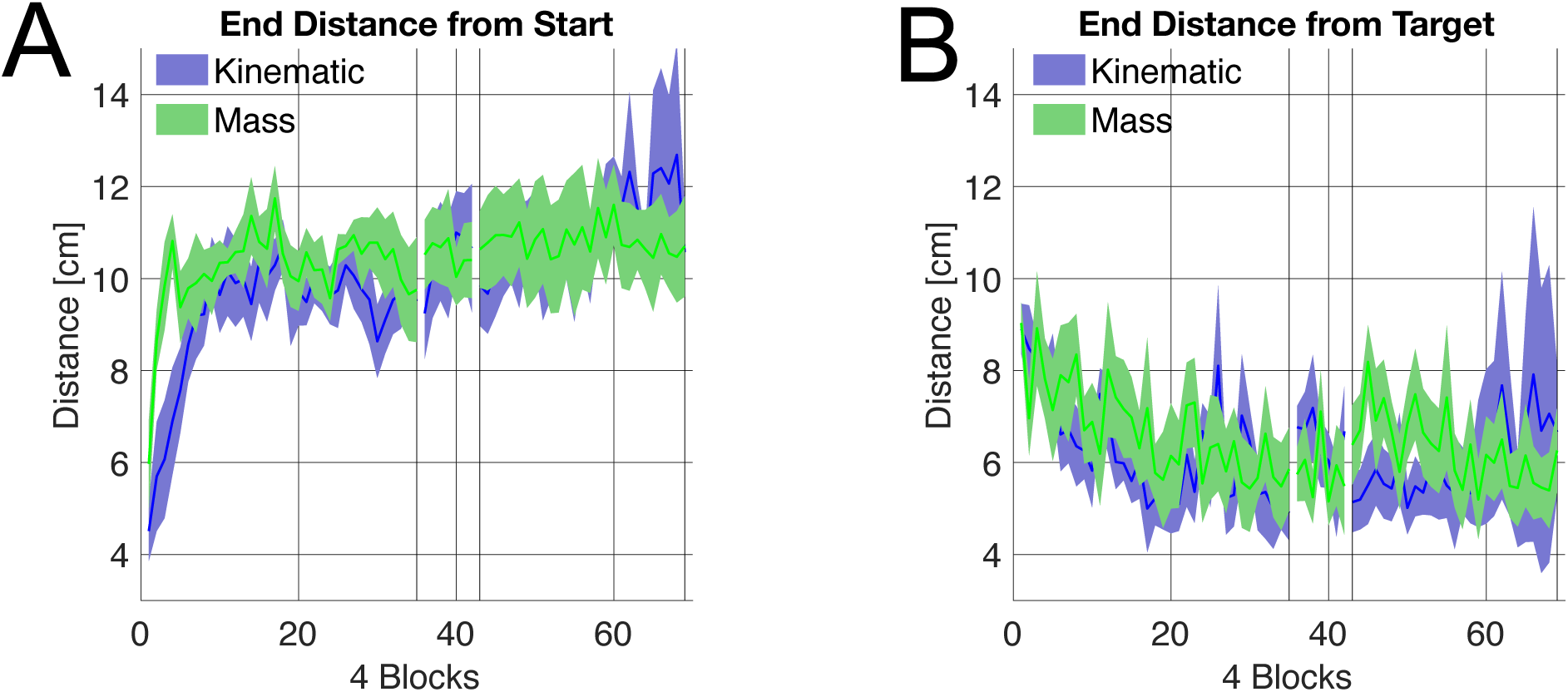
Examination of predictive control. **A:** Deviation from the start point at movement termination. **B:** Deviation from the target point following movement termination.

## Discussion

### Summary

In this study, we examined how learning a novel movement–outcome mapping is shaped by haptic feedback and how such learning influences subsequent adaptation to unpredictable dynamics. We found that participants trained with endpoint dynamics, compared with a kinematic-only control group, showed reduced directional bias during de novo learning and achieved lower residual directional error during subsequent exposure to a velocity-dependent curl-field. Crucially, in both conditions, the presence of robust washout after-effects indicates predictive compensation rather than passive mechanical attenuation. Together, the results indicate that performance in the presence of task-relevant dynamics was associated with altered adaptation to novel perturbations.

Between-condition differences in adaptation could not be explained by movement speed alone: although movement durations differed between conditions, path length did not, and speed-normalized analyses of curl-field adaptation yielded the same pattern of results.

Catch trials (with no visual feedback of the cursor or arm) were used to qualitatively probe feedforward control of the novel mapping. Early in training, catch movements were often misdirected and poorly scaled, consistent with exploratory control. With practice, catch-trial movements became more appropriately scaled in extent and more accurate in direction, indicating improved predictive specification of movement amplitude under feedforward control. These improvements are consistent with the acquisition of a stable feedforward control mapping from bimanual hand inputs to the virtual arm state, rather than reliance on online visual error correction alone.

### Kinematic Task (Experiment 1)

In the Kinematic Task (Experiment 1), participants initially moved their hands sequentially, briefly arresting one hand before advancing the other. With practice, they adopted simultaneous bimanual movements, and extrinsic trajectories progressively straightened. Upon introduction of a curl-field, trajectories initially showed pronounced curvature, but straightened with continued exposure, similar to typical behavior observed in curl-field learning. However, examination of the SMPE suggested that adaptation was incomplete by the end of the curl-field exposure, indicating residual directional bias rather than full predictive compensation for the imposed dynamics.

### Endpoint Mass Task (Experiment 2)

The Endpoint Mass Task (Experiment 2), which added a 2-kg endpoint mass, displayed parallel trends. Early movements again featured independent hand control, but the simulated inertia induced noticeable overshoot. As training progressed, participants quickly adopted simultaneous movements and gradually learned to control the arm to perform point-to-point movements, resulting in straighter extrinsic trajectories with practice. Upon introduction of the curl-field, loopy trajectories were observed, but to a markedly lesser extent than in Experiment 1, and the SMPE was considerably lower. Directional bias was reduced relative to Experiment 1, and signed error approached baseline levels by the end of curl-field exposure, consistent with more complete reduction of directional bias.

A potential concern is that the reduced curl-field-induced curvature in the endpoint-mass condition may not reflect transfer from prior experience alone. In the present design, the simulated endpoint mass was present not only during Day 1 training but also throughout Day 2, including during curl-field exposure and washout. Accordingly, the smaller trajectory deviation observed in Experiment 2, particularly at curl-field onset, may partly reflect the concurrent inertial properties of the task itself. However, both conditions involved the same physical manipulanda and participant limb dynamics. Thus, the comparison was not between an inertial and a non-inertial physical system, but between two groups that differed specifically in the presence or absence of an additional simulated endpoint mass within the task dynamics. The present design therefore does not fully dissociate the effects of prior haptic learning from the ongoing mechanical influence of the endpoint mass during curl-field exposure. Nevertheless, several aspects of the results suggest that the difference between conditions was not exhausted by a trivial effect of movement speed alone. The endpoint-mass condition continued to show lower signed error after normalization by peak velocity, and exponential analysis indicated that the clearest between-condition difference was in asymptotic residual error rather than in learning rate. We therefore interpret the findings cautiously as showing that performing the task in the presence of task-relevant endpoint dynamics was associated with lower residual directional error during curl-field exposure, while acknowledging that the present design cannot determine whether this benefit arose from prior haptic experience, the concurrent endpoint mass on Day 2, or both.

### Learning Rate vs Final Performance

An important observation is that participants were able to adapt to the velocity-dependent curl-field following acquisition of a non-veridical kinematic mapping, with no indication that prior remapping hindered dynamic learning. Although the present study did not include a veridical-control baseline, the observed learning dynamics suggest that the motor system can acquire novel dynamics within an already remapped control space. This flexibility is consistent with the possibility that dynamic learning operates over task-relevant state representations rather than fixed limb-centric coordinates.

We note that differences in adaptation between conditions were not driven by differences in learning rate. Although estimated learning time constants (τ) differed numerically between conditions, permutation-based inference did not provide sufficient evidence for a between-group difference in learning speed. Instead, the primary empirical distinction lay in the final level of residual directional error achieved, with participants in the endpoint-mass condition exhibiting more complete compensation for the curl-field. Together, these findings are consistent with more complete final predictive compensation in the endpoint-mass condition, without evidence of faster adaptation.

### Haptic Feedback and Networks in the Brain

Although the present study is behavioral, the improvement in predictive adaptation is consistent with frameworks in which haptic feedback contributes to internal model formation [37]. Proprioceptive and tactile signals support state estimation and predictive control, processes commonly associated with distributed sensorimotor networks including cerebellar and parietal circuits. Learning in the presence of endpoint dynamics may therefore provide richer state information during de novo acquisition. We restrict our interpretation to the behavioral level and do not infer specific neural mechanisms.

### De Novo Learning

The present task lies at the intersection of two historically distinct literatures: bimanual coordination of shared objects in which each hand contributes to a common task variable [38,39], and the acquisition of complex visuo-mechanical transformations implemented by tools or lever systems [40–42]. In contrast to classical relative-phase paradigms, however, the present mapping does not simply require stabilization of a coordination pattern within a fixed transformation. Instead, participants must learn how two independent control dimensions jointly determine the kinematics of a virtual articulated system.

The relative ease with which participants learned this mapping suggests that the motor system can identify and stabilize novel input–output relationships through exploration and feedback. Early in training, predictive commands were unavailable, and performance relied heavily on exploration of the input space. With practice, variability declined and movements became increasingly coordinated, consistent with progressive stabilization of a novel control mapping.

### Future Work

Future studies could dissociate general force exposure from learning specific dynamic structure by independently manipulating endpoint and velocity-dependent forces. The mechanisms underlying the observed transfer remain to be determined.

## Acknowledgements

We thank John Welsh for invaluable technical support, and David W. Franklin for useful scientific discussion.

## Author Contributions

ISH conceived, designed, and implemented the study. LAH conducted data collection. Data analysis was carried out by ISH. The initial draft of the manuscript was written by ISH. Both authors (ISH and LAH) reviewed and edited the manuscript.

## Data Availability Statement

Data will be made publicly available upon publication via an online repository (details will be provided upon acceptance).

