## Supplementary Materia V4.pdf for "Task-Relevant Haptic Feedback Improves Asymptotic Performance in de novo Arm Control Acquisition" (from another version of this preprint: version 4, 2026-03-02)

for

This Supplementary Information provides full methodological and analytical details that are not included in the main text for brevity. Specifically, it includes:

- The implementation of the mechanical channels used to constrain the left and right handles along the sagittal (y-axis) direction.
- The implementation of the kinematics and dynamics of the two-dimensional (2D) virtual arm.
- The procedures used for exponential fitting of the signed mean path error (SMPE) and absolute mean path error (AMPE) data.
- Supplementary Table S1: Detailed statistical comparisons between phases for Experiment 1.
- Supplementary Table S2: Detailed statistical comparisons between phases for Experiment 2

### Mechanical Channels

The movement of the two vBOT handles was restricted to linear channels aligned with the y-axis, with a length (*chanLen*) of 30 cm. To spatially separate the hands, the left and right channels were offset laterally by ±10 cm, yielding central handle positions at (-10, 0) cm and (+10, 0) cm, respectively.

Simulated mechanical channels constrained hand motion to a single degree of freedom along the y-axis [37], with a wall stiffness of $k_{chan}$ = 4,000 N·m$^{-1}$ and a damping coefficient $b_{chan}$ = 30 N·s·m$^{-1}$. The channel force applied to each handle was given by:

$$\begin{bmatrix} F_{xh} \\ F_{yh} \end{bmatrix} = k_{chan} \begin{bmatrix} -1 & 0 \\ 0 & 0 \end{bmatrix} \begin{bmatrix} x_h - x_o \\ y_h \end{bmatrix} + b_{chan} \begin{bmatrix} -1 & 0 \\ 0 & 0 \end{bmatrix} \begin{bmatrix} \dot{x}_h \\ \dot{y}_h \end{bmatrix} \quad (1)$$

which relates the forces experienced at a given handle, $[F_{xh} \quad F_{yh}]^T$, to its position relative to the channel, $[x_h - x_o \quad y_h]^T$, where $[x_h, y_h]$ denotes the handle position, $x_0$ is the x-location of the channel, and the corresponding handle velocity is $[\dot{x}_h \quad \dot{y}_h]^T$.

### Control of the Simulated Arm

All simulations were executed within the real-time vBOT control loop at 1 kHz. The y-position (y) of each handle was linearly mapped to the corresponding joint angle ($\theta$):

$$\theta = \theta_{min} + (\theta_{max} - \theta_{min}) \frac{\left(dir_{slider} \cdot y + \frac{\text{chanLen}}{2}\right)}{\text{chanLen}} \tag{2}$$

where chanLen is the length of the channel, and $dir_{slider}$ sets the direction in which the slider acts (taking a value of 1 or -1). The y-location of the left handle, $y_{h1}$, was mapped to the shoulder joint angle, $\theta_1$, over the range of $\theta_{min} = 0$ radians and $\theta_{max} = \pi$ radians:

$$\theta_1 = \frac{(15 - y_{h1})}{30} \pi \tag{3}$$

The y-location of the right handle $y_{h2}$ was mapped to a joint angle $\theta_2$, between the range of $\theta_{min} = -\pi$ radians and $\theta_{max} = 0$ radians:

$$\theta_2 = \frac{(15 - y_{h2})}{30} \pi - \pi \tag{4}$$

This mapping defined the reachable workspace shown in Fig. 1E. Differentiating Equations (3) and (4) yields a mapping between the handles' y-velocities and the corresponding joint angular velocities:

$$\begin{bmatrix} \dot{\theta}_1 \\ \dot{\theta}_2 \end{bmatrix} = -\frac{\pi}{30} \begin{bmatrix} \dot{y}_{h1} \\ \dot{y}_{h2} \end{bmatrix} \tag{5}$$

### Simulating Forward Kinematics

Forward kinematics of the 2D arm is required to compute the endpoint position and to determine and display the arm configuration. The simulated arm link lengths $l_1$ and $l_2$, were both 12 cm. Using trigonometric analysis (Fig. 1C), the positions of the elbow and end–effector in Cartesian coordinates can be expressed using the following matrix equations:

$$\begin{bmatrix} x_{elbow} \\ y_{elbow} \end{bmatrix} = \begin{bmatrix} l_1 \cos(\theta_1) \\ l_1 \sin(\theta_1) \end{bmatrix} + \begin{bmatrix} x_{shoulder} \\ y_{shoulder} \end{bmatrix} \tag{6}$$

$$\begin{bmatrix} x_{end} \\ y_{end} \end{bmatrix} = \begin{bmatrix} l_1 \cos(\theta_1) + l_2 \cos(\theta_1 + \theta_2) \\ l_1 \sin(\theta_1) + l_2 \sin(\theta_1 + \theta_2) \end{bmatrix} + \begin{bmatrix} x_{shoulder} \\ y_{shoulder} \end{bmatrix} \tag{7}$$

Differential kinematics of the 2D arm are required to relate handle velocities to endpoint velocities and to map endpoint forces back to forces at the vBOT handles. The Jacobian matrix of the 2D arm is given by:

$$J = \begin{bmatrix} \frac{\partial x_{end}}{\partial \theta_1} & \frac{\partial x_{end}}{\partial \theta_2} \\ \frac{\partial y_{end}}{\partial \theta_1} & \frac{\partial y_{end}}{\partial \theta_2} \end{bmatrix} = \begin{bmatrix} -l_1 \sin(\theta_1) - l_2 \sin(\theta_1 + \theta_2) & -l_2 \sin(\theta_1 + \theta_2) \\ l_1 \cos(\theta_1) + l_2 \cos(\theta_1 + \theta_2) & l_2 \cos(\theta_1 + \theta_2) \end{bmatrix} \tag{8}$$

The relationship between the handle velocities and the extrinsic endpoint velocity is given by the following expression:

$$\begin{bmatrix} \dot{x}_{end} \\ \dot{y}_{end} \end{bmatrix} = J \begin{bmatrix} \dot{\theta}_1 \\ \dot{\theta}_2 \end{bmatrix} = -\frac{\pi}{30} J \begin{bmatrix} \dot{y}_{h1} \\ \dot{y}_{h2} \end{bmatrix} \tag{9}$$

### Task Force Feedback

The arm was modelled as a purely kinematic linkage; all task dynamics were imposed exclusively at the endpoint and mapped to the handles via the Jacobian transpose ($J^T$). The relationship between the virtual arm endpoint force vector $[F_{xend} \quad F_{yend}]^T$ and the joint torque vector $[\tau_1 \quad \tau_2]^T$ is given by:

$$\begin{bmatrix} \tau_1 \\ \tau_2 \end{bmatrix} = J^T \begin{bmatrix} F_{xend} \\ F_{yend} \end{bmatrix} \tag{10}$$

where $J^T$ denotes the transpose of the Jacobian matrix. The torques $[\tau_1 \quad \tau_2]^T$ at the arm joints were mapped to corresponding y-direction forces, $[F_{h1} \quad F_{h2}]^T$, applied at the vBOT handles by accounting for the mechanical advantage relating handle displacement to joint rotational angles. This mapping yields the following conversion formula:

$$\begin{bmatrix} F_{h1} \\ F_{h2} \end{bmatrix} = dir_{slider} \frac{(\theta_{max} - \theta_{min})}{\text{chanLen}} \begin{bmatrix} \tau_1 \\ \tau_2 \end{bmatrix} \tag{11}$$

Substituting the expression for the joint torques and the corresponding numerical values yields the final expression for the force along the y-axis at the handle in terms of endpoint force:

$$\begin{bmatrix} F_{h1} \\ F_{h2} \end{bmatrix} = -\frac{\pi}{30} J^T \begin{bmatrix} F_{xend} \\ F_{yend} \end{bmatrix} \tag{12}$$

### Endpoint Dynamics Simulation

In the null-field condition, when the arm endpoint was unloaded, no endpoint forces were generated or experienced at the control handles during arm movement. When endpoint dynamics were introduced, namely, a viscous curl-field, an added mass, or a combination of both, resulting endpoint forces were mapped back to the handles and perceived by participants as they moved the arm through the workspace.

### Endpoint Curl-field Simulation

In curl-field endpoint trials, a velocity-dependent curl-field [12] was implemented by relating arm endpoint velocity to the endpoint force:

$$\begin{bmatrix} F_{xend} \\ F_{yend} \end{bmatrix} = b_{curl} \begin{bmatrix} 0 & -1 \\ 1 & 0 \end{bmatrix} \begin{bmatrix} \dot{x}_{end} \\ \dot{y}_{end} \end{bmatrix} \tag{13}$$

The curl-field constant $b_{curl}$ was set to ±16 N·s·$m^{-1}$, where the sign determined the direction of the curl-field (clockwise or counterclockwise). In each experiment, the curl-field direction was counterbalanced across the eight participants to minimize directional bias. Each participant experienced only one curl-field direction.

The endpoint velocity used in Equation (13) was calculated from the handle velocities using Equation (9). The corresponding forces experienced along the y-direction at the handles were then calculated using Equation (12). The total two-dimensional force applied to the vBOT handles was computed as the sum of the channel force component and the force feedback component arising from the curl-field.

### Simulating Endpoint Mass

Endpoint mass dynamics were simulated by attaching a virtual point mass to the arm endpoint via a stiff linear spring (k_mass = 2000 N·$m^{-1}$). Viscous damping was also included in the simulation to improve stability. The mass was initialized at the endpoint position with zero velocity and the spring at rest length; initial acceleration and applied force were set to zero. The arm and endpoint mass were simulated within the 1 kHz real-time vBOT control loop. At each timestep, the mass position was updated using forward Euler integration:

$$\begin{bmatrix} x_m \\ y_m \end{bmatrix} = \begin{bmatrix} x_m \\ y_m \end{bmatrix} + \begin{bmatrix} v_{xm} \\ v_{ym} \end{bmatrix} \Delta t \tag{14}$$

Where $\Delta t$ is the timestep since the previous update (nominally 1 ms). Acceleration of the mass was estimated using the equation:

$$\begin{bmatrix} a_{mx} \\ a_{my} \end{bmatrix} = \frac{1}{m} \begin{bmatrix} F_{xm} \\ F_{ym} \end{bmatrix} \tag{15}$$

Where m is the mass at the endpoint load (2 kg) and $[F_{xm} \quad F_{ym}]^T$ is the force acting on the mass. Euler integration of the acceleration of the mass was used to calculate endpoint mass velocity:

$$\begin{bmatrix} v_{xm} \\ v_{ym} \end{bmatrix} = \begin{bmatrix} v_{xm} \\ v_{ym} \end{bmatrix} + \begin{bmatrix} a_{xm} \\ a_{ym} \end{bmatrix} \Delta t \tag{16}$$

The force acting on the mass was then computed from extension of the spring, with spring constant $k_{mass}$ = 2000 N·m⁻¹, connecting the endpoint mass to the endpoint of the arm:

$$\begin{bmatrix} F_{xm} \\ F_{ym} \end{bmatrix} = -k_{mass} \left( \begin{bmatrix} x_m \\ y_m \end{bmatrix} - \begin{bmatrix} x_{end} \\ y_{end} \end{bmatrix} \right) \tag{17}$$

An additional viscous damping term was applied to the endpoint to improve stability:

$$\begin{bmatrix} F_{xdamp} \\ F_{ydamp} \end{bmatrix} = -b_{end} \begin{bmatrix} \dot{x}_{end} \\ \dot{y}_{end} \end{bmatrix} \tag{18}$$

Where damping viscosity $b_{end}$ = 10 N·s·m⁻¹. The mass force (Equation 17) and damping force (Equation 18) were summed, and the corresponding forces experienced in the y-direction at the handles were then calculated using Equation (12). The total two-dimensional force applied to the vBOT handles corresponded to the sum of the channel force component and the endpoint force-feedback components.

### Simultaneous Endpoint Mass and Curl-field

In trials with both an endpoint mass and a curl-field present, endpoint mass and curl-field components were computed as described previously. The vBOT applied an overall two-dimensional force to its handles by summing the force-feedback components from the curl-field and the endpoint mass, together with the channel forces that restrict handle movement to the y-direction.

### Learning-curve Analysis

#### Exponential model fitting

Exponential fitting was performed as a descriptive summary of learning dynamics and to complement block-level comparisons reported in the main text. Learning analyses were restricted to SMPE during curl-field exposure, as unsigned metrics (e.g., AMPE) do not isolate predictive compensation and may decrease through non-adaptive strategies such as co-contraction.

Learning dynamics were quantified by fitting an exponential model to block-pair averaged performance,

$$y(t) = Ae^{-t/\tau} + C \tag{19}$$

where $A$ represents the deviation from asymptote at $t = 0$ (such that $y(0) = A + C$), $\tau$ is the learning time constant, and $C$ is the asymptotic performance level. Here, $t$indexes block pairs (each data point representing the mean of 32 curl-field trials, excluding catch trials).

Exponential models were fit to block-pair data pooled across participants within each condition. Individual participant fits were explored but were unstable due to variability across block pairs and were not used for inference.

Group-level parameter differences were evaluated using a participant-label permutation procedure. Participants from both conditions were pooled and randomly reassigned to two pseudo-groups of the same sizes as the original groups, without replacement. For each permutation, exponential models were fit to pooled block-pair data within each permuted group, and parameter differences were recorded. Two-sided p-values were computed as the proportion of permuted differences whose absolute magnitude exceeded the observed difference. This approach avoids distributional assumptions and directly tests the null hypothesis of label exchangeability. Parameter estimates reported for each condition were obtained from the observed group assignments; permutation was used solely to construct the null distribution of between-group parameter differences.

### Supplementary Table S1: Detailed statistical comparisons between phases for Experiment 1

Table S1: Repeated measures ANOVAs for within condition test in Experiment 1

| Metric | | From 1st to final null | Decrease Final null to first curl | Reduction First curl to last curl | Last curl to washout |
|---|---|---|---|---|---|
| SMPE | $F(4,28) = 26.061$<br>$p < 0.001$<br>$\omega^2 = 0.617$ | $p_{bonf} = 1.000$ | **$p_{bonf} < 0.001$** | **$p_{bonf} = 0.049$** | **$p_{bonf} = 0.002$** |

| | | | | | |
|---|---|---|---|---|---|
| AMPE | $F(4,28) = 11.366$<br>$p < 0.001$<br>$\omega^2 = 0.463$ | **$p_{bonf} = 0.027$** | **$p_{bonf} < 0.001$** | **$p_{bonf} = 0.043$** | **$p_{bonf} = 0.047$** |
| Movement Duration | $F(4,28) = 30.213$<br>$p < 0.001$<br>$\omega^2 = 0.728$ | **$p_{bonf} = 0.006$** | **$p_{bonf} = 0.009$** | **$p_{bonf} = 0.029$** | $p_{bonf} = 0.639$ |
| Extrinsic Path Length | $F(4,28) = 11.269$<br>$p < 0.001$<br>$\omega^2 = 0.466$ | $p_{bonf} = 0.061$ | **$p_{bonf} < 0.001$** | **$p_{bonf} = 0.008$** | **$p_{bonf} = 0.044$** |

### Supplementary Table S2: Detailed statistical comparisons between phases for Experiment 2

Table S2: Repeated measures ANOVAs for within condition test in Experiment 2

| Metric | | From 1st to final null | Decrease Final null to first curl | Reduction First curl to last curl | Last curl to washout |
|---|---|---|---|---|---|
| SMPE | $F(4,28) = 14.013$<br>$p < 0.001$<br>$\omega^2 = 0.504$ | $p_{bonf} = 1.000$ | **$p_{bonf} = 0.003$** | **$p_{bonf} = 0.013$** | **$p_{bonf} < 0.001$** |
| AMPE | $F(4,28) = 23.674$<br>$p < 0.001$<br>$\omega^2 = 0.629$ | **$p_{bonf} = 0.014$** | **$p_{bonf} < 0.001$** | **$p_{bonf} = 0.004$** | **$p_{bonf} = 0.004$** |
| Movement Duration | $F(4,28) = 166.581$<br>$p < 0.001$<br>$\omega^2 = 0.923$ | **$p_{bonf} < 0.001$** | **$p_{bonf} = 0.016$** | **$p_{bonf} < 0.001$** | **$p_{bonf} = 0.031$** |
| Extrinsic Path Length | $F(4,28) = 23.558$<br>$p < 0.001$<br>$\omega^2 = 0.647$ | **$p_{bonf} = 0.013$** | **$p_{bonf} < 0.001$** | **$p_{bonf} = 0.009$** | **$p_{bonf} = 0.020$** |
